# Expression of a transcript encoding for cytosolic renin in the diseased human heart

**DOI:** 10.1101/2020.04.02.021592

**Authors:** Jaroslaw Sczodrok, Philipp Lutze, Doreen Staar, Marcus Dörr, Jörg Peters

## Abstract

Secretory renin promotes hypertrophy, apoptosis, necrosis and fibrosis through angiotensin generation. It has been claimed that local expression of renin contributes to the deleterious effects of the renin-angiotensin system in the heart. Besides the classic renin transcript (renin-a), encoding for secretory renin, a putative brain-specific (renin-b) and a putative lung-specific (renin-c) transcript may exist in human. In contrast to secretory renin, renin-b cannot be secreted, remains within the cytosol and is imported into mitochondria. In contradiction to renin-a, renin-b exerts cardioprotective effects in hearts and in cardiac cells of the rat under ischemia related conditions. To date, available data on cardiac renin expression remain inconsistent. Nobody has yet investigated which renin transcripts are expressed in the human heart. We systematically analyzed the levels of renin transcripts using specific and sensitive nested reverse-transcriptase polymerase chain reactions (RT-PCR) in human ventricular biopsies obtained from patients with heart diseases. In the 33 biopsies available, neither the expression of classic renin-a, nor of the alternative renin-c was detected. In contrast, the renin-b transcript, which was previously classified as brain-specific, was found in 11/33 ventricular biopsies. Our data exclude the expression of secretory renin and indicate that local expression of cytosolic renin but not of secretory renin can play a functional role in the human heart.

## Introduction

Renin is a secretory glycoprotein that, after secretion into the extracellular space, generates angiotensin (ANG) I from its only known substrate, angiotensinogen [1]. ANG I is further cleaved to ANG II, the effector peptide of the renin-angiotensin system (RAS) that increases oxidative stress and exerts pro-inflammatory effects [2, 3]. In the heart, ANG II promotes hypertrophy, apoptosis, necrosis, fibrosis, myocardial remodeling and hence cardiac failure [4, 5]. Correspondingly, inhibitors of the RAS are among the most potent drugs in the treatment of hypertension and cardiac failure, markedly increasing the life span of patients [6].

In addition to the classic renin transcript, alternative renin transcripts have been identified in several species including mice, rats and humans [exon(2-9), exon(1A-9), renin-b, renin-c] [7–9]. These alternative transcripts lack exon 1. Exon 1 encodes for the signal sequence required for a co-translational transport to the endoplasmatic reticulum and thus for the sorting of the renin protein to a secretory pathway. As a consequence the translated renin protein cannot be secreted, remains in the cytosol and is taken up by mitochondria [8, 10]. All alternative renin transcripts are translated into a truncated prorenin with use of the first in-frame UTG in exon 2. The protein lacks the pre-fragment and the first 15 amino acids of the conventional prorenin [7–9].

The alternative transcripts were described to be exclusively expressed in brain (renin-b) or lung (renin-c) [7, 9]. However, in rat we detected expression in many extrarenal tissues including the adrenal gland and heart [11]. In the rat heart, the expression of cytosolic renin but not of secretory renin was increased after myocardial infarction. Furthermore, cells overexpressing cytosolic renin are more stable against necrotic cell death [10, 11], particularly under ischemic conditions such as glucose depletion and anoxia combined with glucose depletion [12, 13]. Also, overexpression of renin-b in the heart reduced the infarct size in Langendorff preparations ex vivo [12]. Therefore, we ask whether or not - and which - alternative renin transcripts are expressed in the human heart and analyzed biopsies of patients with coronary heart disease (CHD), hypertensive heart disease (HHD) and cardiomyopathies (CM).

## Methods

### Study design

The present study comprised 33 patients with suspected dilated cardiomyopathy (DCM) and/or myocarditis. In all subjects, an endomyocardial biopsy (EMB) was taken for diagnostic reasons in accordance to the position statement of the European Society of Cardiology Working Group on Myocardial and Pericardial Diseases [14]. For assessment of myocardial virology and immunohistochemistry were obtained from the right ventricular (RV) septum. Acute infectious diseases, cancer, chronic alcoholism, postpartum cardiomyopathy, coronary heart disease or heart failure due to other known origins (e.g. primary valvular disease) were excluded. Acute myocarditis was excluded by histopathological analysis of EMBs in accordance with Dallas criteria [15] Familial DCM was excluded according to patient information collected by questionnaire. With written consent of each patient EMBs that were stored for clinical purposes as reserve material for immunohistochemistry and virology, but which were not needed any more for routine diagnostics were available for protein extraction. The investigation conforms to the principles of the Declaration of Helsinki and the protocol was approved by the Ethics Committee of the University of Greifswald, Germany (Reg. number III UV 34/04).

#### RNA Isolation and RT-PCR

PolyA-RNA from heart, brain and lung of adult men was purchased by AMS Biotechnology (Europe) Ltd, Abingdon Oxon OX14 4RX, UK). Human kidney total RNA was used as reference [16]. RNA isolation of human left ventricular biopsies was performed with TRIzol (Invitrogen/ThermoFisher Scientific Inc., Waltham, MA, USA). The cardiac tissue samples (0.3 – 5.4 mg) were ground in a sterile, cooled porcelain mortar under liquid nitrogen. 200 – 300 μl TRIzol reagent were added, homogenized for 10 s, then incubated for 5 min. Cell debris were removed by centrifugation (500 g, 5 min, 4 °C). 50 μl chloroform were added to the supernatants and carefully inverted. After an incubation time of 3 min, the samples were centrifuged (12000 g, 15 min, 4 °C). To eliminate lipids, proteins and DNA the upper aqueous phase was carefully removed while not disturbing the interphase. RNA was precipitated with the addition of 150 μl isopropanol. After 15 min incubation at 4 °C the samples were centrifuged (12000 g, 20 min, 4 °C), washed twice with 300 μl ethanol (12000 g, 15 min, 4 °C), then dried on air. The pellets were dissolved in 20 μl TE-buffer (10 mM Tris, 1 mM EDTA), samples were stored at −70 °C. Subsequently the isolated or purchased RNAs were used for RT-PCR with MMLV reverse transcriptase (Ambion Inc., Austin, TX, USA), following manufacturer’s instructions to obtain cDNA for further PCR analyses.

#### PCR protocols

Nested PCR was performed in order to amplify transcripts of renin-a, -b and -c, respectively (n= 4 – 12). PCR parameters were chosen as following: 94 °C initial temperature for 3 min, 35 cycles with each 94 °C denaturation step for 30 sec, 56 °C annealing step for 20 sec, 68 °C elongation step for 40 sec, followed by a final elongation step at 72 °C for 5 min (SureCycler 8000, Agilent technologies, Santa Clara, CA, USA). In order to amplify different transcripts via nested PCR from the *renin* gene, primers (5’ → 3’) were designed as outer and inner pairs.

The products of the outer PCR were used as templates for the amplification with inner primers. Amplification was carried out for renin-a using the forward-outer primer (fwd-out): GACTGCTGCTGCTGCTCTG together with the reverse-outer primer (rev-out): TGTAGAGACGGCTGCACTTG, followed by forward-inner primer: (fwd-in): TACCTTTGGTCTCCCGACAG together with the reverse-inner primer (rev-in) AAAGACGACTTTGAAGGTCTGG. For, renin-b we used the fwd-out CCACACAACAGCAAGTAAGCTG together with the rev-out GATGCCAATCTCGCCATAGT, followed by fwd-in: CAACAGCAAGTAAGCTGAGAGG together with the rev-in CGCCATAGTACTGGGTGTCC. For renin-c we used the fwd-out CTCAGGGACTCCACCACTG together with the rev-out ATTCTCGGAATCTCTGTTGTAG, and fwd-in CAGCACTTTCTATTTTTGCTTCC together with the rev-in ATTCCACCCACGGTGATGAT. For Tyrosine 3-Monooxygenase/Tryptophan 5 Monooxygenase Activation Protein, Zeta (YWHAZ) we used fwd GCAACCAACACATCCTATCAGAC together with rev CCTCCTTCTCCTGCTTCAGC. Inner PCR’s yielded bands of following sizes: renin-a, 250 bp; renin-b, 188 bp; renin-c, 443 bp and YWHAZ, 249 bp (Tab. I; online supplement).

#### Agarose electrophoresis

The PCR products were separated with pre-stained (Roti-GelStain; Carl Roth GmbH + Co. KG, Karlsruhe, Germany) 1.5% agarose gel using purple gel loading dye and 2-Log DNA ladder (both New England Biolabs, Ipswich, MA, USA) at 80 V and 85 mA for 2h. Visualization of the bands was performed using ChemiDoc XRS+ imaging system with Image Lab software (Bio-Rad Laboratories Inc., Hercules, CA, USA).

#### Subcloning and sequencing

Distinct bands of correct size were cut from 1.5% agarose gel and isolated using illustra GFX PCR and Gel Band Purification Kit (GE Healthcare, Chicago, IL, USA), following manufacturer’s protocol. After ligation into pGEM-T Easy Vector (Promega Corp., Madison, WI, USA), the plasmids were transformed into *E. coli* Stellar cells (Clontech/Takara Bio USA Inc., Mountain View, CA, USA). Cells were grown to an OD600 of about 0.5 and afterwards harvested. Plasmids were isolated using PureYield Plasmid Miniprep System (Promega) and sanger sequencing was performed (GATC Biotech AG, Konstanz, Germany).

## Results

For this study we had access to 33 left ventricular biopsies of patients with different heart diseases (28 males, 5 females, 39 – 73 years; for additional information see Tab. II, online supplement). The patients were subdivided into 4 groups according to their predominant disease: Hypertensive heart disease, HHD (n=10), coronary heart diseases, CHD (n=8), cardiomyopathy, CM (n=13) and other heart-related diseases (pericardial effusion and respiratory global insufficiency, each n=1). Since no tissue from healthy human hearts was available for obvious reasons, we had to perform our analyses with purchased RNA of human men in order to verify the specificity of the PCR protocols as well as the expression profile of the three renin transcripts. Therefore we analyzed transcripts of renin-a, -b and -c in human polyA-RNA from heart, brain and lung as well as total RNA from kidney obtained from [16] (Fig.1). Although we never detected renin-a in the total RNA preparation of the biopsies (see below), renin-a was detected in the cardiac polyA-RNA. Since polyA-RNA has about a hundred-fold higher mRNA levels than total RNA, we expect, that this finding of minor amounts of renin-a is irrelevant. Also, even with all given carefulness an unwanted contamination cannot be excluded entirely. Amplification of renin-a from human kidney using the nested PCR approach resulted in two bands for unknown reasons, one of them at the correct size of 250 bp. However, when performing the inner PCR with human kidney cDNA without previous outer amplification, only this single distinct band appears. Furthermore, analysis of the outer PCR’s without further inner amplification showed that the transcript for renin-a only appears in the kidney and not in the heart (data not shown).

**Fig. 1.**
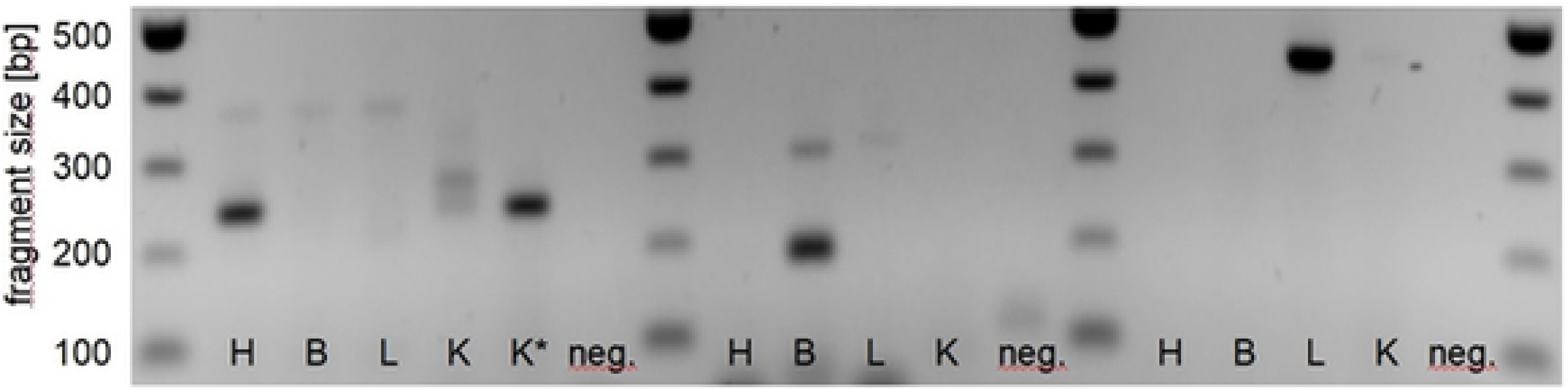
Transcripts of renin-a (250 bp, left), renin-b (188 bp, middle) and renin-c (443 bp, right) in control RNAs. H = heart, B = brain, L = lung, K = kidney, K* = kidney (inner amplification shown without outer PCR), neg. = negative control (H_2_O).

As previously described, transcripts of renin-b were detected in polyA-RNA from human brain, while transcripts of renin-c were solely detected in human lung.

For the systemic analyses of human biopsies, we confirmed RNA isolation by amplifying *YWHAZ* as a commonly used housekeeper gene. Bands with the correct size were visible in all of our samples (Fig. 5).

Nested PCR with primers for the transcript of the secretory renin-a showed distinct bands in human kidney and also in the heart. PolyA-RNA controls of brain and lung did not show any particular bands of correct size. With the described PCR setup the transcript of the secretory renin-a remained undetectable in all of our 33 biopsies (Fig. 2). Modulating PCR parameters (template concentration, annealing temperature, touchdown-PCR approaches, number of cycles; n=6) resulted in the same overall picture, where no renin-a transcripts were found in human biopsies.

**Fig. 2.**
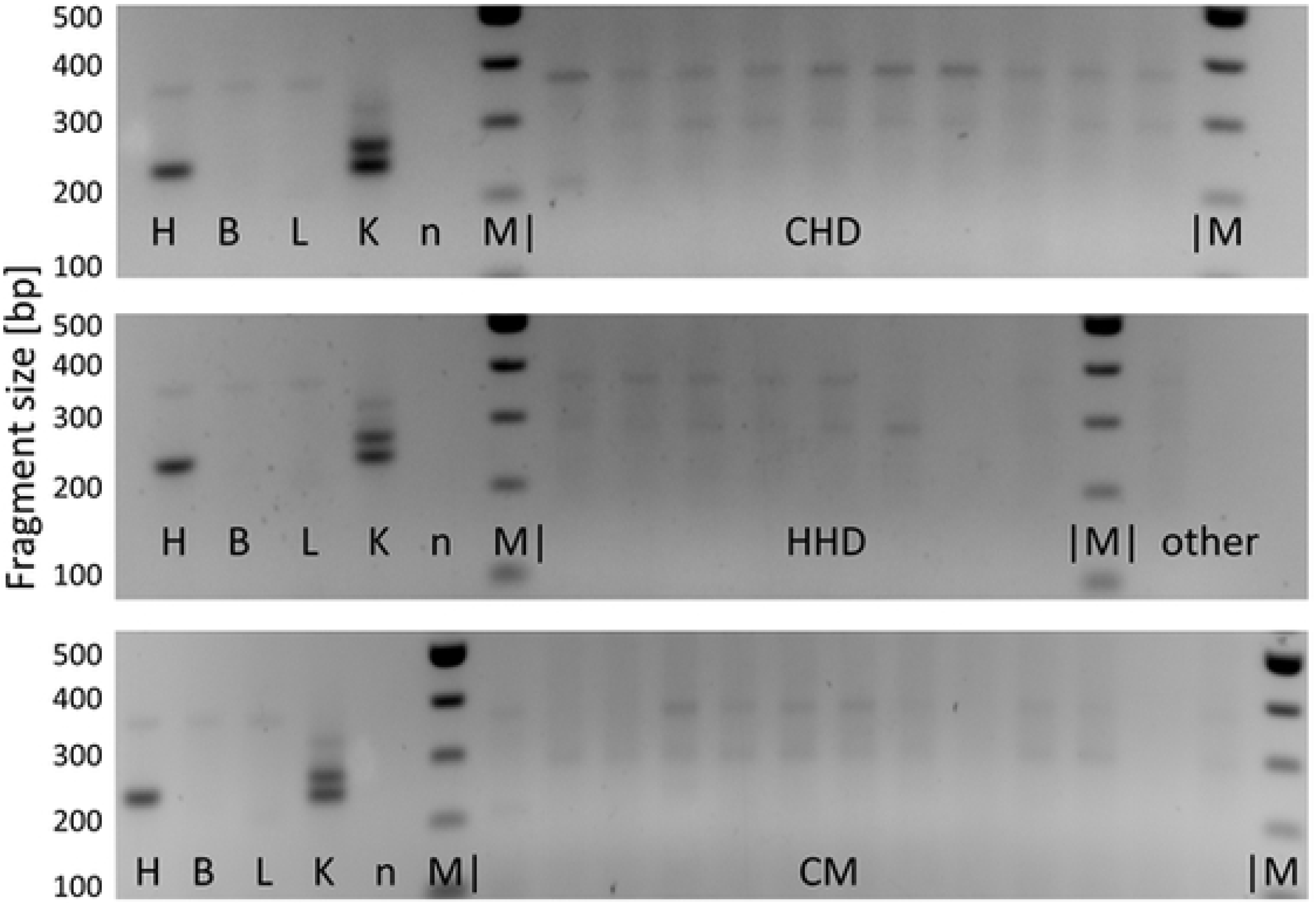
Transcripts of renin-a (250 bp) in human cardiac ventricular biopsies; groups: CHD, (upper panel), HHD and other (middle panel), CM (lower panel). H = heart, B = brain, L = lung, K = kidney, n = negative control (H_2_O), M = DNA marker

The lung-specific transcript for the non-secretory renin-c was not detected in any of the 33 biopsies using highly specific nested PCR protocols (Fig. 3). This remains true for all other PCR parameters we have tested (n=4). Thus, it can be seen as not expressed in human heart under pathological conditions.

**Fig. 3.**
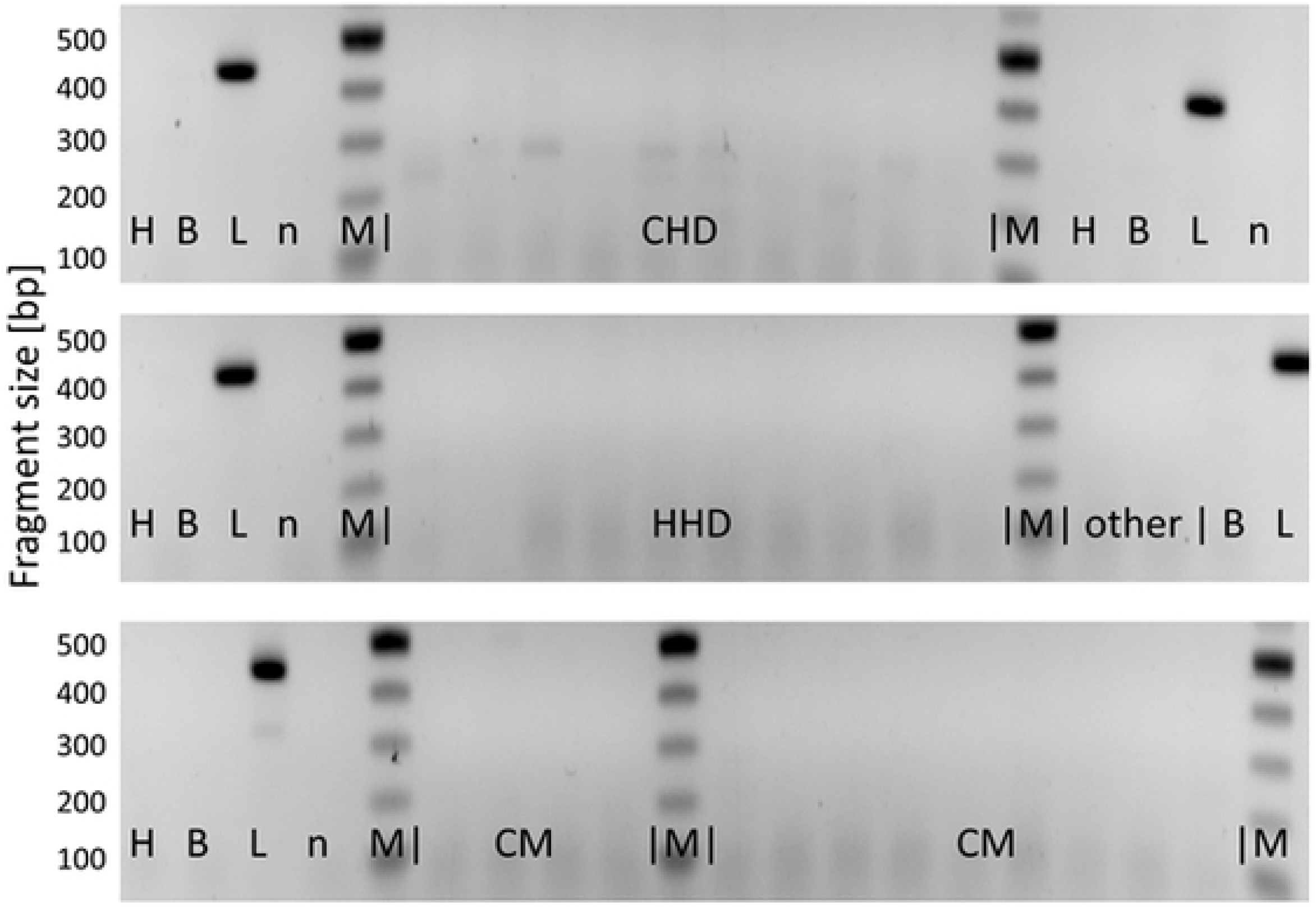
Transcripts of renin-c (443 bp) in human cardiac ventricular biopsies; groups: CHD, (upper panel), HHD and other (middle panel), CM (lower panel). H = heart, B = brain, L = lung, n = negative control (H_2_O), M = DNA marker

On the other hand, the transcript of the brain-specific non-secretory renin-b was detected in the human brain and also a small amount was found in polyA-RNA from human heart. In the biopsies of our patients, we have identified transcripts of renin-b in 11/33 cases: 3/10 ventricular biopsies from patients with CHD, in 3/8 biopsies from patients with HHD, in 3/13 biopsies from patients with CM and in 2/2 patients with other heart-related diseases (Fig. 4). Randomly chosen samples which resulted in renin-b bands of correct size were isolated to verify the presence of renin-b by sanger sequencing (n=6; Tab. III, online supplement). In summary, we provide evidence for the expression exclusively of renin-b in human left ventricular biopsies under pathological conditions in 11/33 biopsies within all groups. Transcripts of renin-a and renin-c could not be detected.

**Fig. 4.**
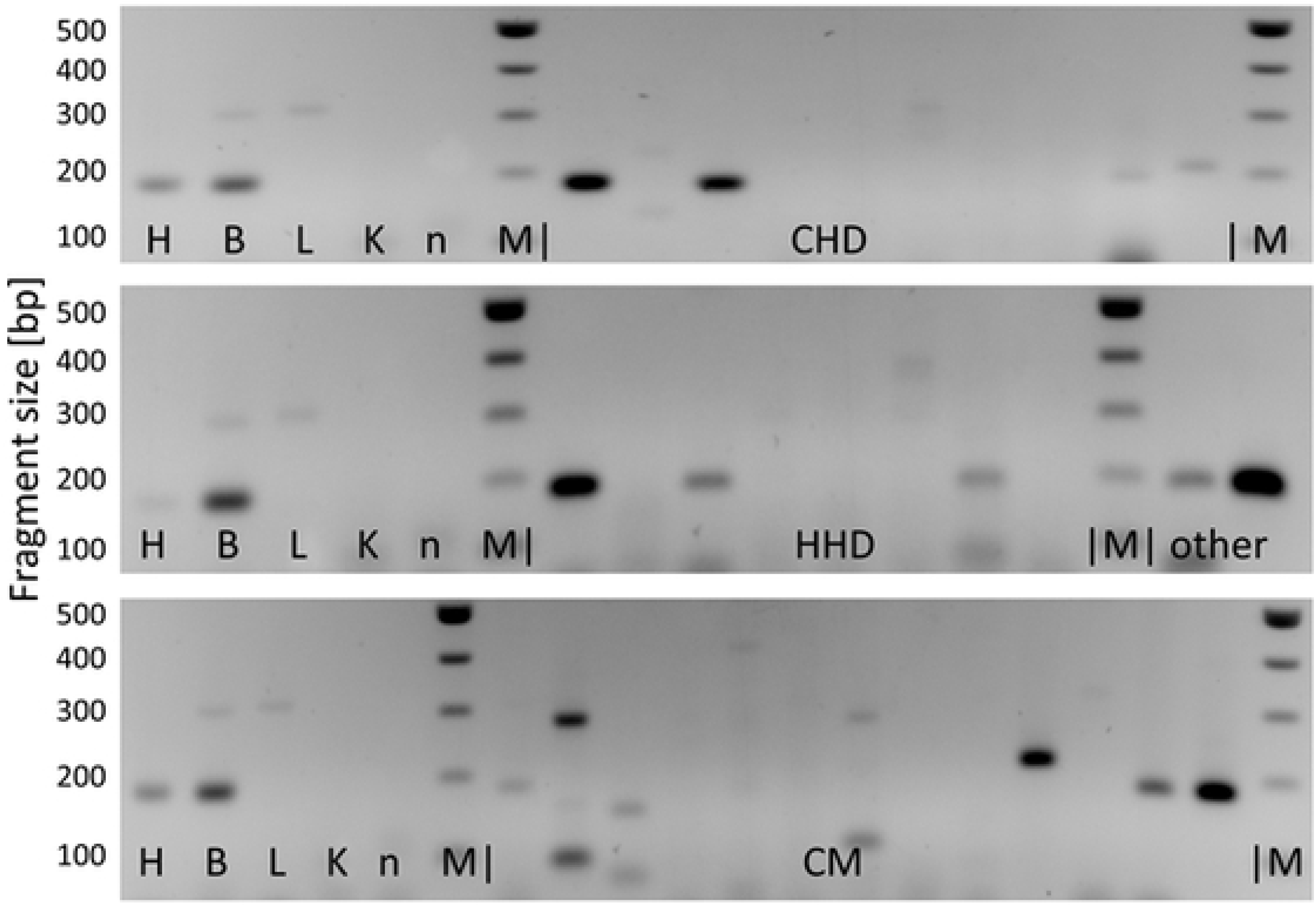
Transcripts of renin-b (188 bp) in human cardiac ventricular biopsies; groups: CHD, (upper panel), HHD and other (middle panel), CM (lower panel). H = heart, B = brain, L = lung, K = kidney, n = negative control (H_2_O), M = DNA marker

**Fig. 5.**
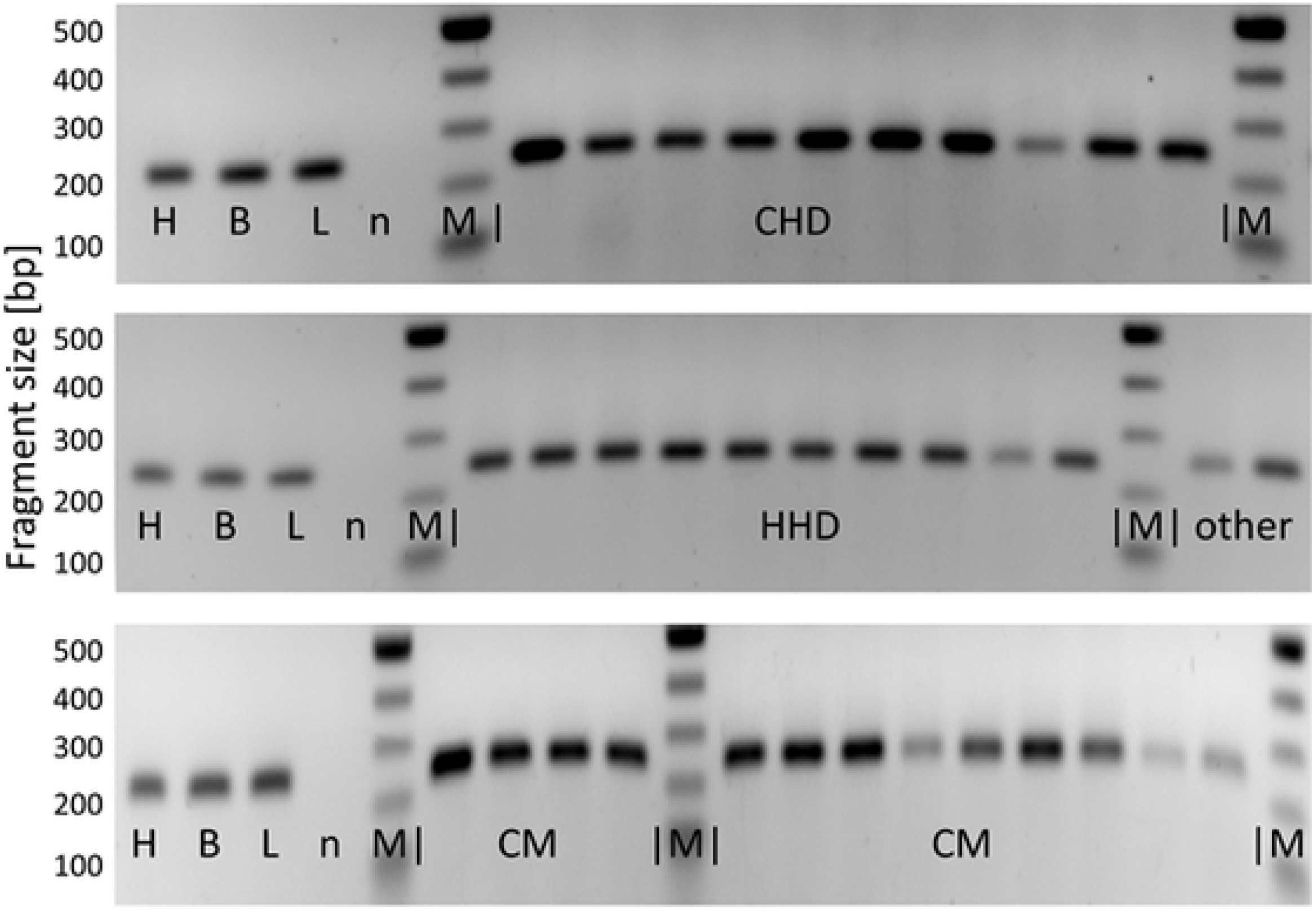
Transcripts of YWHAZ (249 bp) in human cardiac ventricular biopsies; groups: CHD, (upper panel), HHD and other (middle panel), CM (lower panel). H = heart, B = brain, L = lung, n = negative control (H_2_O), M = DNA marker

## Discussion

There is no doubt that renin-angiotensin systems reside locally within the heart [17, 18]. Even the existence of intracellular acting RASs has been postulated [for review see: [19]. However, one would like to know whether or not renin is just taken up or locally expressed in the heart. Uptake of renin would rather reflect the activity of the circulating RAS, determined by the regulation of kidney renin release. On the contrary, local expression of renin in the heart may well be regulated by independent factors, giving rise to (partially) kidney-independent renin systems. For the rat it was demonstrated that the renin enzymatic activity of the heart is derived exclusively from the circulation [20, 21]. This was concluded from the observation that after bilateral nephrectomy renin activity became undetectable.

Results on the expression of renin within the heart have been inconsistent. Although in some studies the presence of renin mRNA was demonstrated [22–26]; others failed to detect any renin transcript under non-stimulated conditions [27–30]. Several studies reported a marked increase in renin expression after myocardial infarction [25, 31], during cardiac hypertrophy [24] and after stretch of cardiomyocytes [32, 33]. The discrepancies regarding cardiac renin expression can be explained by several factors. First, there may be species specific differences; second, the expression profile is age dependent (embryonal, neonatal, adult); third, the expression depends on a specific pathophysiological context (ischemia, stretch), and, finally, there are different renin transcripts derived from the same gene which are differentially expressed. These transcripts are identical form exon 2 to 10 and cannot be differentiated when using nonspecific primers. On the other hand, when using a primer in exon 1, the presences of alternative renin transcripts are overlooked. This argument not only explains some of the discrepancies, but also provides a molecular basis for the existence of a cytosolic-mitochondrial renin (-angiotensin) system in the heart with different modes of action.

So far, most studies on renin expression have been done in rat or mice. Interestingly, in autopsies of human left ventricles renin mRNA was not found in healthy hearts, but was present in subjects with heart diseases [34]. However, in that study non-specific primers spanning exons eight to ten were used, which did not differentiate between the various renin transcripts. Given the fact that different renin transcripts are expressed we now wanted to know more precisely which of the possible renin transcripts are expressed in the human heart. Our data strongly support the hypothesis that circulating secretory renin is solely taken up from the circulation in human, since we were not able to detect any transcripts in the heart encoding for secretory renin, even with most sensitive approaches. In contrast we consistently detected renin-b in human hearts and thus conclude that this transcript is not brain-specific as previously suggested. These results are in agreement with our data in the rat [11].

Since we had no chance to obtain cardiac biopsies from healthy individuals we were not able to answer the question whether or not this renin-b transcript is indeed upregulated in human cardiac disease. Therefore, we could only take advantage of animal models. In this respect we previously demonstrated that in ischemic heart disease specifically the transcript encoding for cytosolic renin was upregulated in rats [11] and that in cardiac rat cells ischemia-related conditions such as glucose depletion or combined anoxia with glucose depletion increased expression of cytosolic renin [12, 13, 35].

In opposition to the harmful effect of secretory renin, renin-b is even protective, particularly under ischemia-related conditions. Thus in the rat we observed less necrosis in long term cultures [10] and less necrotic cell death after glucose depletion or anoxia combined with glucose depletion in cardiac cells overexpressing renin-b [12, 13], as well as a markedly reduced infarct size after ischemia in the Langendorff heart preparations [12] in hearts obtained from transgenic rats [36] overexpressing renin-b. It is important to note that inhibitors of the RAS exclusively inhibit the consequences of the harmful secretory renin enzymatic activity but may not have any effect on the protective renin-b, since at least some of the effects of the latter does not involve angiotensin generation [12]. On the other hand, mitochondrial and nuclear angiotensin receptors have been detected and ANG II indeed exerts effects on isolated mitochondria and nuclei [37, 38]-Chedar), supporting the model of cytosolic angiotensin generation. However, again the effects of ANG II then were always harmful, unless AT1 receptors were blocked or absent. The existence of renin transcripts encoding for cytosolic nevertheless provides an important requirement for cytosolic or mitochondrial generation of angiotensins, which may be protective via AT2 receptors. Thus, intracellular angiotensin generation may still well be considered. Other intracellular interaction partners for cytosolic renin may mediate angiotensin-independent effects of renin-b. One possible target is the cytosolic RnBP that has already been shown to interact with renin in vitro particularly under conditions with depletion of high-energy nucleotides [39, 40]. Such conditions may prevail during ischemia. RnBP acts as an epimerase, interconverting N-acetyl-D-glucosamine and N-acetyl-D-mannosamine. This enzymatic activity of RnBP is inhibited by renin [41]. By regulating the levels of N-acetyl-D-glucosamine, RnBP may affect the glycosylation and sialylation of many proteins, including transcription factors and thus modulate their activities.

In the rat we observed a specific upregulation of renin-b expression under various ischemia-related conditions. Thus, glucose depletion in cardiac cells in vitro and ligation of left coronary arteries in rats in vivo increased transcript levels of renin-b but not of secretory renin [11]. Therefore, we suggested that renin-b transcript levels are increased also in human heart diseases. We do not have direct evidence about the regulation of renin-b in human heart diseases, because we did not have access to healthy biopsies. Thus it must remain unclear whether or not renin-b expression is induced under pathological conditions such as CHD, HHD or CM. However, there are indirect arguments for an induction of renin-b expression in the human heart under pathological conditions. Previous studies on human heart autopsies did not detect renin mRNA in 15 out of 15 ventricles of hearts from patients without cardiac disease, but renin mRNA was present in 2 out of 2 hearts from patients with coronary heart disease and 1 out of 1 heart with hypertrophy [34]. We now have analyzed a larger number of hearts of living patients with coronary artery disease, hypertensive heart disease and cardiomyopathies and differentiated between transcripts encoding for secretory and cytosolic renin isoforms. Taken together it is likely that renin-b mRNA levels but not secretory renin levels increase under pathological conditions in the heart. Inhibitors of the renin-angiotensin system will not be able to interfere with this rather protective renin-b in the heart, targeting only the deleterious secretory renin.

## Author contribution statement

J.S. designed the experiments and methods, performed the analysis, analyzed the data and wrote the manuscript. P.L designed the methods and analyzed the data. D.S. analyzed the data and participated in writing the manuscript. M.D. provided the biopsies, analyzed the data and wrote the manuscript. J.P designed the experiments and methods, analyzed the data and wrote the manuscript.

## Acknowledgements

We thank B. Sturm for excellent technical assistance.

## Data sharing statement

There are no data available to share exept those presented in the figures and supplement

